# A potential link between gambling addiction severity and central dopamine levels – evidence from spontaneous eye blink rates in gamblers and controls

**DOI:** 10.1101/236109

**Authors:** David Mathar, Antonius Wiehler, Karima Chakroun, Dominique Goltz, Jan Peters

## Abstract

Accumulating evidence points at similarities between substance use disorders and pathological gambling on the behavioral and neural level. In substance addiction, dysregulation of striatal dopamine transmission has been consistently revealed. Due to the neurotoxicity of stimulating substances, it is still debated if this constitutes mainly a consequence of recurrent substance abuse or a vulnerability marker for addiction disorders. For gambling addiction, no clear association with striatal dopamine levels has been unveiled so far. With its presumably negligible dopaminergic toxicity, possible differences in striatal dopamine transmission in gambling addiction might therefore constitute a vulnerability marker.

Spontaneous eye blink rate (sEBR) is controversially discussed as a potential proxy measure for striatal dopamine levels. Here we examined sEBR in 21 male problem gamblers and 20 healthy control participants. In addition, participants completed a screening questionnaire for overall psychopathology and self-reported measures of alcohol and nicotine consumption. We found no significant difference in sEBR between gamblers and controls. However, in gamblers, sEBR was negatively associated with addiction severity and positively associated with psychopathology. A final exploratory analysis revealed that healthy controls with low sEBR displayed higher alcohol and nicotine consumption than healthy participants with high sEBR. Although the association between dopamine transmission and sEBR is still debated, our findings reveal that sEBR is sensitive to inter-individual differences in addiction severity in problem gamblers.

## 1 Introduction

Placing something valuable at risk with the hope of gaining something of greater value is a popular recreational activity among adults, and is referred to as gambling (Potenza, 2006). Approximately five percent encounter problematic gambling behavior and around one percent display pathological gambling (Schaffer et al., 1999; Petry et al., 2005). Gambling disorder has been classified as an addiction disorder in the Diagnostic and Statistical Manual of Mental Disorders (5th ed.; DSM–5; American Psychiatric Association, 2013). This classification is based on accumulating evidence revealing similarities between pathological gambling and substance use disorders (SUDs) in both behavioral observations and underlying neural mechanisms (Potenza et al., 2008; Frascella et al., 2010; Genauck et al., 2017).

A consistent finding in substance-based addiction is the dysregulation of dopaminergic function within striatal target sites. Patients suffering from SUDs show reduced dopamine D2-receptor availability in striatum (Volkow et al., 1993, 2001; Heinz et al., 2004; Martinez et al., 2004; Fehr et al., 2008), constituting a candidate risk factor for the development of SUDs. In monkeys and rodents, attenuated striatal dopamine D2-receptor availability prior to drug exposure is related to a more rapid acquisition of drug self-administration (Nader et al., 2006; Dalley et al., 2007). Reduced D2-receptor modulation of corticostriatal pathways may translate into a higher risk of escalating drug abuse via several possible mechanisms. First, critical behavioral markers such as impulsivity (e.g. attenuated inhibitory control, steeper devaluation of delayed rewards) and heightened reward-related cue-reactivity are associated with lower D2-receptor availability in both humans and rodents (Dalley et al., 2007; Oswald et al. 2007; Weber et al., 2016). Second, D2-receptor function is also tightly coupled to learning from negative outcomes (Frank et al., 2007; Mathar et al., 2017), a process recently shown to be impaired in cocaine addicts compared with healthy controls (Parvaz, et al., 2015).

In pathological gambling, the evidence for an association with striatal D2-receptor availability is mixed. Earlier studies reported reduced dopamine levels in cerebrospinal fluid of gambling addicts (Bergh et al., 1997; Meyer et al., 2004). In line with this observation, dopaminergic medication in Parkinson’s disease patients can induce problem gambling as a side-effect (Voon et al., 2009). In contrast, more recent positron-emission-tomography (PET) studies quantifying striatal D2/3-receptor availability observed no differences between gambling addicts and controls (Linnet et al., 2011; Clark et al., 2012; Boileau et al., 2013). However, in pathological gamblers, striatal D2/3-receptor availability was associated with mood related impulsivity and gambling severity (Clark et al., 2012; Boileau et al., 2013). In addition, van Holst et al. (2017) found increased striatal dopamine synthesis capacity, as assessed with [^18^F]fluoro-levo-dihydroxyphenylalanine ([^18^F]DOPA) PET, in gamblers compared with controls.

Spontaneous eye blink rate (sEBR) is discussed as a potential non-invasive proxy measure of striatal dopamine transmission (Karson et al., 1983). Initial evidence linking sEBRs to dopamine transmission came from observations in several neurological and psychiatric disorders that relate to alterations in central dopamine regulation such as Parkinson’s disease, schizophrenia and psychosis (Freed et al, 1980; Karson et al., 1982; Mackert et al., 1991; Colzato et al., 2009). In line with reduced D2/3-receptor availability, cocaine users show reduced sEBRs compared with healthy controls (Colzato et al., 2008). Several pharmacological studies observed that sEBR is reduced following dopamine antagonist administration, and elevated after dopamine agonist administration (Blin et al. 1990; Elsworth et al., 1991; Kleven and Koek 1996; Kaminer et al., 2011). Complementing these findings, a PET study in monkeys found a strong positive correlation between sEBR and D2-receptor availability in ventral and parts of dorsal striatum (Groman et al., 2014). Consistent with the idea that sEBR measures trait-like differences in dopaminergic functioning, it has good reliability (Cronbach’s Alpha: .79-.85, see Kruis et al., 2016). However, two recent studies report opposing findings. Dang et al. (2017) found no significant correlation between sEBR and D2/3-receptor availability in midbrain and striatum in humans. In addition, they observed no significant impact of bromocriptine, a dopamine agonist, on participants’ sEBRs. Noteworthy, their subject sample was quite heterogeneous regarding age and body weight, and between 3 and 32 months separated subjects’ sEBR assessment from PET imaging. In a preprint, Sescousse et al. (2017) report a negative correlation between sEBR and striatal dopamine synthesis capacity in a mixed sample of healthy controls and gamblers. Thus, the association between sEBR and striatal dopamine still needs further consideration.

Here, we utilize sEBR to examine potentially dopamine D2-receptor modulated group differences between gambling addicts and healthy control participants. We hypothesize that gamblers show reduced sEBRs compared with healthy controls. As psychopathology is known to be related to aberrant central dopamine levels (Maas et al., 1993; Gray, 1995) and recently was shown to correlate with sEBR (Colzato et al., 2009), we utilize the SCL-90-R questionnaire (Derogatis & Unger, 2010) as a screening test, and to control for overall psychopathology.

## 2 Methods

### 2.1 Subjects

21 male problem gamblers, and 20 healthy male control subjects participated in this study. All gamblers reported regular gambling, and suffered from losing money while gambling. 12 gamblers fulfilled DSM-IV criteria (#criteria>=5) of pathological gambling. Gamblers and healthy controls were matched for age, educational background, socioeconomic status, alcohol and nicotine consumption. Severity of gambling addiction was assessed by the ‘Kurzfragebogen zum Glücksspielverhalten’ (KFG, Petry et al., 1996), and a German adaptation of the South Oaks Gambling Screen (SOGS, Lesieur & Blume, 1986). Both questionnaires are validated screening tools for quantifying gambling addiction, and show good reliability (Cronbach’s Alpha: KFG: .79; SOGS: .97). Comprehensive demographic information is provided in Table 1. Participants were recruited via advertisements on local internet bulletin boards. Prior to enrollment in the study, phone interviews were conducted, and only gamblers who reported gambling on a regular basis, suffered from monetary loss, and fulfilled at least one of the DSM-IV criteria of pathological gambling were invited to participate. Gamblers were mainly engaged in slot machine gambling (67%), and sports betting (57%). A fraction also pursued (online) poker (14%), and roulette (14%). Eligible participants were screened to exclude a history of neurological or psychiatric disorders, current medication, and substance abuse other than nicotine and alcohol. All study procedures were approved by the local Institutional Review Board (Hamburg Board of Physicians), and participants provided informed written consent prior to their participation. Participants received 10 EUR per hour as a compensation for participation.

**Table 1.**

|  | healthy controls | gamblers | F/U | p |
| --- | --- | --- | --- | --- |
| n | 20 | 21 | - | - |
| age | 26.4 ± 6.39 (19-45) | 26.0 ± 6.66 (18-42) | F=.04 | 0.85 |
| income | 1028.15 ± 575.05<br>(0-2000) | 1375.1 ± 819.51<br>(300-2700) | F=2.44 | 0.13 |
| YOE | 11.75 ± 1.37 (9-14) | 11.71 ± 1.82 (9-15) | F=.01 | 0.94 |
| BDI-II | 8.7 ± 8.3 (0-28) | 15.1 ± 11.87 (2-42) | F=3.96 | 0.05 |
| GSI | .32 ± .34 (0-1.1) | .73 ± .62 (1-2.46) | F=6.7 | 0.01 |
| DSM-5 | .1 ± .38 (0-1) | 5.1 ± 2.28 (1-8) | U=1 | 1.1*10<sup>-8</sup> |
| KFG | 1.45 ± 4.07 (0-18) | 25.29 ± 14.54 (6-54) | U=9 | 7.6*10<sup>-8</sup> |
| SOGS | .4 ± 1.0 (0-4) | 8.48 ± 4.61 (3-17) | U=5.5 | 3.7*10<sup>-8</sup> |
| GA | -.77 ± .21 (-.85-.08) | .73 ± .86 (-.38-2.3) | U=217 | 8.2*10<sup>-8</sup> |
| AUDIT | 6.75 ± 4.8 (0-15) | 5.95 ± 6.98 (0-23) | F=.43 | 0.52 |
| #cigarettes | 9.25 ± 8.75 (0-30) | 5.95 ± 6.76 (0-19) | U=167.5 | 0.25 |
| sEBR | 14.39 ± 4.98 (4.8-22.8) | 15.03 ± 5.59 (4.8-22.8) | F=0.14 | 0.7 |

Sample description [mean ± standard deviation (min-max)]: Gamblers did not differ regarding age, income, years of education (YOE), alcohol (AUDIT) and nicotine consumption (#cigarettes), and eye blink rate (sEBR). Gamblers displayed higher gambling addiction (DSM-5, KFG, SOGS, sum of z-scores of KFG & SOGS (GA)), higher psychoticism (GSI), and a tendency for more depressive symptoms (BDI-II). Tests for group differences were based on ANOVA (F) for normally distributed variables, and Mann-Whitney U Tests otherwise.

### 2.2 General procedure

Participants entered the lab in the afternoon around 2 pm. After they gave written informed consent, participants started with a five minutes sEBR assessment. Notably, sEBRs are stable throughout the day and rise in the evening (Barbato et al., 2000). Subsequently, they completed a questionnaire battery. On a separate day, participants performed a reward learning task in an fMRI setting. These data will be reported elsewhere.

### 2.3 Psychological assessment

Besides the assessment of gambling severity, participants completed the Symptom Check-List-90-R (SCL-90-R, Derogatis & Unger, 2010) that constitutes a screening tool for capturing current psychological pathology. As depressive symptoms are a common co-morbidity in pathological gambling (Petry et al., 2005) participants also completed the Beck Depression Inventory (BDI-II, Beck et al., 1961).

In addition, we quantified nicotine consumption as self-reported number of cigarettes smoked per day, and alcohol use was measured utilizing the Alcohol Use Disorders Identification Test (AUDIT, Saunders et al., 1993).

### 2.4 SEBR assessment

Spontaneous eye blink rates (SEBRs) were assessed via electromyography (EMG), utilizing a MP100 system running under the software Acqknowledge 3.9.1 (Biopac Systems, Goleta, California). Data was recorded via three Ag-AgCL electrodes with a sample rate of 1000Hz, and an online bandpass filter of 28-500Hz. One reference electrode was placed on the middle of the participant’s forehead and two electrodes were fixed below the lower lash line of the left eye, one of the electrodes centrically and the other one 2-3mm in peripheral proximity. Participants sat in front of a computer screen and were instructed to move as little as possible while staring at a fixation cross at approximately 0.5m distance for 5 minutes. They were not explicitly told that sEBR was assessed.

Individual sEBRs were computed using Matlab 2012b (MathWorks, Natick, MA) via the ‘*findpeaks*’ function in a sliding window approach of 10 seconds to the acquired raw data. Blinks were defined as peaks exceeding the data’s mean within the moving window by six standard deviations. SEBRs were then calculated as average number of blinks per minute.

### 2.5 Statistical analyses

All reported results were computed with PASW-SPSS-Statistics 17.0 (IBM Corporation, Somers, NY, USA). We utilized an ANOVA model to assess differences in sEBR between pathological gamblers and healthy controls. Stepwise (backward elimination) multiple regression analysis was used to test whether sEBR is related to gambling addiction severity in gamblers. Gambling addiction severity was computed as the mean of the two z-standardized gambling questionnaire scores (SOGS + KFG). Notably, results were similar if only one score was used in the regression model. To control for overall psychopathology, the global severity index (GSI) of the SCL-90-R served as an additional regressor. We further controlled for age-related effects. In a second regression model, we controlled for individual depressive symptomatology via including individual BDI-II scores instead of gamblers’ GSI scores. We did not include both regressors in one model, as both variables were highly correlated (R^2^=.78, p=7.44*10^-8^).

In addition, we ran an exploratory analysis to test an association between sEBR and substance use in healthy controls. An individual substance use score was calculated as the sum of the z-standardized AUDIT questionnaire score and the z-standardized number of cigarettes smoked per day. Controls were separated into a low and a high sEBR group via a median split to test if the low sEBR group consumed more alcohol and /or nicotine according to the substance use score than the high sEBR group. Additionally, we computed a correlation analysis between sEBR and GSI scores in healthy controls.

Gaussianity, heteroscedasticity and absence of multicollinearity were tested for the respective analyses.

## 3 Results

SEBRs did not differ significantly between healthy controls (HC) and gamblers (G) (mean +/- SEM HC: 14.6 ± 1.1; G: 14.4 ± 1.4; F(1,39)=.02, p=.89, Fig 1a). If only pathological gamblers were included (DSM-IV >=5), sEBRs were still similar to healthy controls (PG: 13.4 ± 1.7; F(1,30)=.41, p=.53). In gamblers however, stepwise multiple regression analysis (adj. R^2^=.31, F(2,18)=5.43, p=.01, Table 2, final model) revealed that sEBR was negatively correlated with gambling addiction severity (mean [95% CI]: β=-.53 [-5.61 -.69], p=.02, Fig 1b) and positively correlated with overall psychopathology (GSI) (β=.54 [.1 .78], p=.01, Fig 1c). Age showed no significant impact on sEBR and thus was removed from the model.

**Figure 1.**
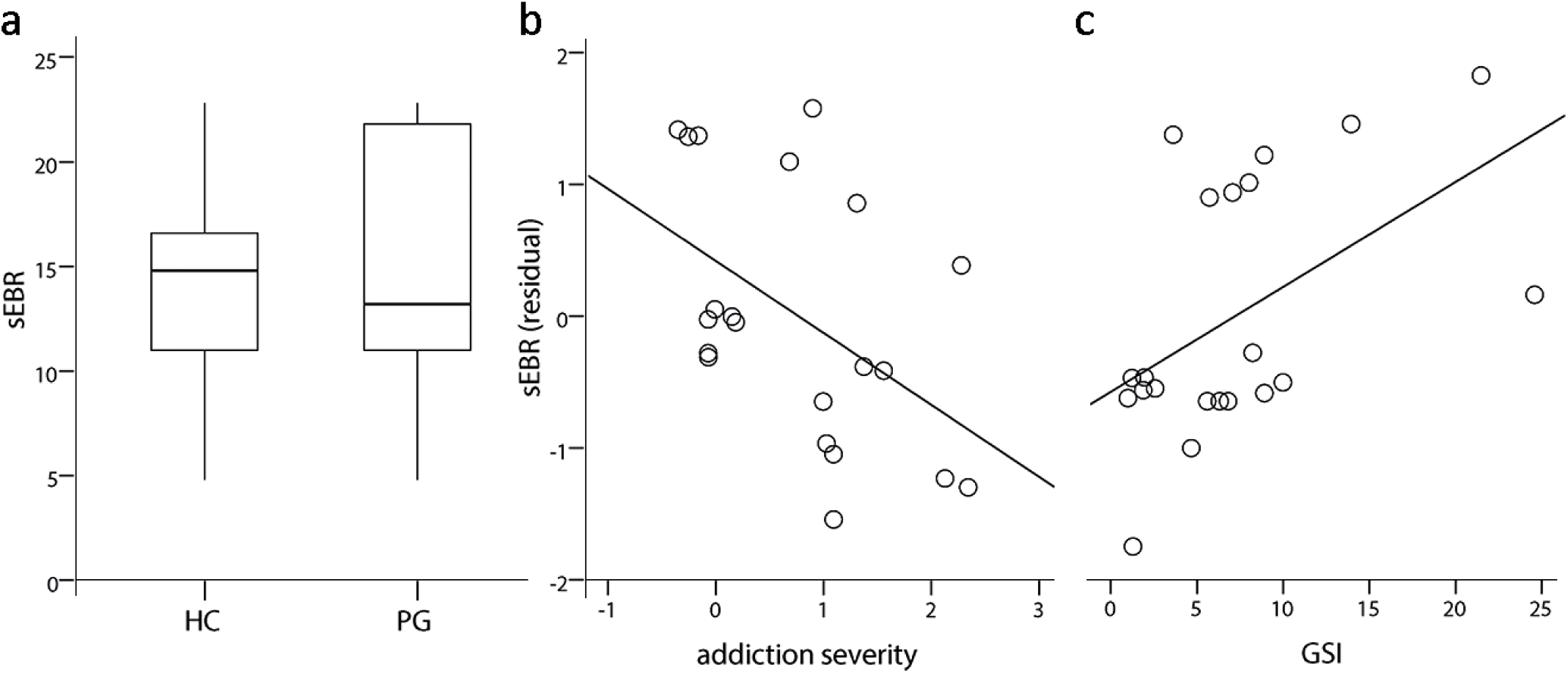
Notes: (a) SEBR did not differ in problem gamblers (PG) compared with healthy controls (HC). (b) Multiple regression analysis revealed that addiction severity in pathological gamblers negatively correlated with sEBR. (c) Manifestation strength of psychotic symptoms (GSI) was positively associated with sEBR in gamblers.

**Table 2.**

|  | final model | initial model | alternative model |
| --- | --- | --- | --- |
| GA | $\beta = -.53$ [-5.61 -.69]<br>p=.02 | $\beta = -.54$ [-5.88 -.45]<br>p=.03 | $\beta = -.45$ [-5.68 .09]<br>p=.06 |
| GSI | $\beta = .54$ [.1 .78], p=.01 | $\beta = .55$ [.05 .85]<br>p=.03 | - |
| age | - | $\beta = .008$ [-.35 .36]<br>p=.97 | - |
| BDI-II | - | - | $\beta = .31$ [-.08 .34]<br>p=.2 |
| model statistics | adj. $R^2 = .31$,<br>F(2,18)=5.43, p=.01 | adj. $R^2 = .27$<br>F(3,17)=3.42, p=.04 | adj. $R^2 = .11$<br>F(2,18)=2.22, p=.14 |

Computed regression models: Gambling addiction (GA) and psychopathology (GSI) predicted sEBR in gamblers according to stepwise regression (final model). Age (initial model) was no significant predictor of sEBR. Depressive symptoms (BDI-II) instead of GSI scores did not explain significant variance in gamblers' sEBR (alternative model). Values in brackets represent 95% confidence intervals.

Individual GSI scores were highly correlated with BDI-II scores (adj. R^2^=.78, F(1,19)=71.31, p=7.44*10^-8^). To exclude that the impact of psychopathology on gamblers’ sEBRs was exclusively driven by depressive symptoms, we computed another regression model that included BDI-II scores instead of participants GSI scores and addiction severity. This model did not explain a significant amount of variance of individual sEBRs (adj. R^2^=.09, F(2,18)=2.0, p=.16, Table 2, alternative model) as BDI-II scores did not predict sEBRs significantly (β=.31 [-.09 .38], p=.2).

In an exploratory analysis (see *Statistical analyses*), we tested whether sEBR in healthy controls was associated with substance use (alcohol and nicotine use). For this purpose, we computed a compound score of alcohol and nicotine consumption and separated our control participants into a low sEBR and a high sEBR group according to a median split.

Control participants with a low sEBR showed higher nicotine and alcohol consumption than participants with a high sEBR (F(1,18)=5.92, η^2^=.25, p=.03, Fig 2). In contrast to gamblers, GSI scores were not correlated with sEBRs in control participants (R^2^=.13, p=.11).

**Figure 2.**
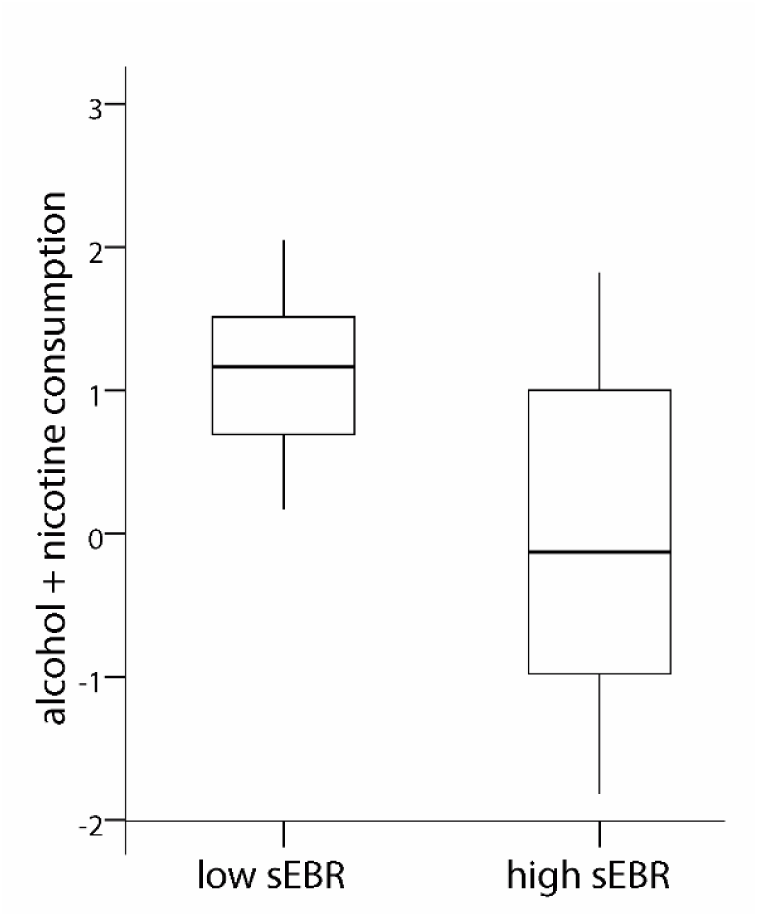
Notes: Healthy controls with low sEBR consumed more alcohol and nicotine (z-standardized) than control participants with high sEBR.

## 4 Discussion

We utilized spontaneous eye blink rate (sEBR) to test for potentially striatal dopamine modulated group differences between problem gamblers and matched healthy control participants. We observed no differences between gamblers and controls. However, in gamblers, addiction severity was negatively associated with sEBR, suggesting a potential modulatory effect of striatal dopamine levels on escalation speed of gambling behavior. Overall psychopathology, as assessed with the SCL-90-R questionnaire, was positively linked to sEBR in gamblers. Additional exploratory analysis revealed that in healthy controls, sEBR was negatively associated with alcohol and nicotine use.

Our observation that gamblers displayed similar sEBR compared with matched controls supports findings from direct assessments of striatal D2/3 receptor availability recently conducted in gambling addicts and controls (Clark et al., 2012, Boileau et al., 2013).

Thus, reduced striatal D2/3 availability may not constitute a risk factor for gambling addiction, in contrast to substance-based addictions where reduced D2/3 availability is a consistent finding in both animals (Nader et al., 2006, Dalley et al. 2007), and humans (Volkow et al., 2001; Martinez et al., 2009). Likewise, reduced sEBR has been related to recreational cocaine use in humans (Colzato et al., 2008). Drug consumption presumably causes a higher subsequent increase in striatal dopamine levels than engagement in gambling. Thus, gambling may not suffice to compensate for reduced striatal dopamine as it is assumed for drug use. Noteworthy, recurrent abuse of stimulating substances also leads to a decrease in striatal D2/3 receptor availability (Nader et al., 2006; Volkow et al., 1999; Groman et al., 2012). This aggravates the differentiation between cause and consequence regarding altered dopamine signaling and addiction disorders at least in cross-sectional studies in humans.

We found that in problem and pathological gamblers, gambling addiction severity was negatively associated with sEBR as a potential marker of striatal dopamine levels. This is in line with a recent observation of Boileau et al. (2013). They found that in pathological gamblers D3 receptor availability, as assessed via [^11^C]-(+)-propyl-hexahydro-naphtho-oxazin (PHNO) binding potential in the substantia nigra, positively correlated with gambling addiction severity. As [^11^C]-(+)-PHNO is highly sensitive to fluctuations in extra synaptic dopamine levels (Willeit et al., 2006), this may at least partly translate into a negative association between central dopamine levels and addiction severity. Taken together, this indicates that reduced central dopamine levels may not serve as a vulnerability marker for the development of gambling addiction similar to SUD, but instead may exacerbate addictive behavioral patterns in gamblers when gambling habits are already established.

Overall psychopathology correlated positively with sEBR as a potential indicator of striatal dopamine levels in gamblers, but not in healthy controls. Psychological aberrations such as psychoticism has previously been related to heightened sEBR (Colzato et al., 2009; Barbato et al., 2012) and reduced striatal D2 receptor availability (Gray et al., 1994). The consistent association of schizophrenia and heightened striatal dopamine function (Gray et al., 1991; Di Forti et al., 2007; Eyles et al., 2012) together with a continuum model of psychosis (Johns and van Os, 2001) further support the hypothesis that a psychosis-prone personality has a dopaminergic basis (Ettinger et al., 2013). In healthy controls, the range of the GSI score was significantly lower than in problem and pathological gamblers with a maximum score of 1.1 compared to 2.46, respectively. This complements earlier observations of an association between mental health disorders and pathological gambling (Cunningham-Williams et al., 1998; Grant et al., 2005; Desai & Potenza, 2009), and may at least partly explain the absence of a correlation between GSI scores and sEBR in control participants.

In an exploratory analysis, we found that healthy controls displaying relatively low sEBR (i.e. potentially low striatal dopamine levels) consumed more alcohol and nicotine than participants with relatively high sEBR. This is in accordance with findings from PET studies showing reduced D2/3 receptor availability in alcohol and nicotine addiction (Hietala et al., 1994; Heinz et al., 2005; Brody et al., 2004; Martinez et al., 2005; Volkow et al., 2007; Fehr et al., 2008). Notably, alcohol and nicotine consumption both increase extracellular dopamine in striatum (Boileau et al., 2003; Brody et al., 2004; Barrett et al., 2004) that likely leads to a downregulation of striatal dopamine signaling following chronic intake (Hietala et al. 1994; Martinez et al., 2005; Fehr et al., 2008). Thus, this finding may at least partly reflect a consequence of recurrent consumption.

### 4.1 Limitations

Several limitations of the present study need to be acknowledged.

First, the assumption of a positive correlation between sEBR and dopamine levels in striatum in humans is still under debate. In a recent preprint, Sescousse et al., (2017) found a negative relation between sEBR and dopamine synthesis capacity as assessed with [ 18F]DOPA PET in a mixed sample of gamblers and control participants. Dang et al. (2017) found no significant correlation between D2/3 receptor availability and sEBR in humans. Notably, their sample of 20 subjects was quite heterogeneous in age (20-50y) and body weight (<60 – 120kg). Furthermore, PET imaging and sEBR assessment were separated on average by 17 months (3-32months). Hence, more work is needed to clarify the relationship between sEBR and dopamine function in humans.

Second, this was a cross-sectional study. Thus, we cannot exclude that higher sEBR in gamblers exhibiting more severe gambling may be a consequence of gambling history potentially due to adaptations in the dopaminergic system similar to observations in substance-based addiction.

Third, only male participants were tested, limiting the generalization of the findings.

Fourth, our sample of gamblers consisted predominantly of slot-machine and sports betting gamblers, thus limiting our conclusions to this particular subgroup of gamblers.

Finally, the sample size was insufficient to examine potential differences between gambling subtypes, that have for example been proposed in the “pathways model” (Blaszczynski & Nower, 2002).

### 4.2 Conclusion

In light of the potential link between sEBR and striatal dopamine levels, our findings in gamblers and healthy controls indicate that attenuated striatal dopamine activity is not necessarily a risk factor for developing gambling addiction as postulated for substance-based addictions. Rather, attenuated striatal dopamine might aggravate engagement in gambling in problematic gamblers. One endophenotype of lower striatal dopamine function is heightened impulsivity (Oswald et al., 2007; Lee et al., 2009; Buckholtz et al., 2010). Pathological gamblers suffering from relatively low striatal dopamine function may be prone to a more rapid escalation of addictive behavior through attenuated cognitive control, heightened cue-reactivity and/or steeper delay discounting (Peters et al., 2011; Miedl et al., 2012, 2014; Weber et al., 2016).

Given that sEBR proves to be an affordable and easily obtainable measure with a putative link to dopaminergic function, it might be worthwhile to explore its applicability in clinical practice. For example, it would be of interest to explore sEBR as a potential predictor of treatment outcome in addiction and/or examine interindividual differences in sEBR changes posttreatment.

## Author Notes

J.P. and A.W. designed research. A.W. performed research. D.M. and A.W. analyzed data. K.C. and D.G. contributed analytical tools. All authors contributed to writing of the paper. This research was funded by Deutsche Forschungsgemeinschaft (PE 1627/5-1 to J.P.).

The authors declare no conflict of interest.

